# The Portable Microhaplotype Object and Tools

**DOI:** 10.64898/2025.12.10.693568

**Authors:** Nicholas J. Hathaway, Kathryn Murie, Maxwell Murphy, Alfred Simkin, Jorge Amaya-Romero, Alfred Hubbard, Jessica Briggs, Andrés Aranda-Díaz, Angela M. Early, Amy Wesolowski, Daniel E. Neafsey, Jeffrey A. Bailey, Bryan Greenhouse

## Abstract

**Structured Abstract:** *Motivation:* The rapid increase in the generation of targeted sequencing data offers immense potential for research, medicine, and public health, however the lack of an established standard for these data has led to disparate solutions for data storage. A widely accepted standard is essential for data sharing, reuse, and the coordinated development of interoperable analysis tools.

*Results:* We propose the Portable Microhaplotype Object (PMO), a standardized format for efficiently and losslessly storing phased targeted sequencing data (microhaplotypes). The PMO format is JSON-based, allowing efficient, relational storage of genetic data together with relevant metadata to minimize orphaned data. The format includes required fields and a curated set of optional fields leveraging established ontologies. To facilitate ease of use, we developed pmotools-python, an open-source package for creating, manipulating, and exporting PMO data into common formats. Additionally, we provide a simple web-based app to quickly create PMO files from tabular inputs, making the format accessible to a wide variety of users. Example datasets from *Plasmodium, Anopheles, Escherichia coli*, and *Staphylococcus aureus* demonstrate the broad applicability of the approach. PMO will streamline data sharing, foster interoperability, and accelerate the development of harmonized analysis tools.

*Availability and implementation:* The Portable Microhaplotype Object (PMO) project, including the ontology specification, software tools, example datasets, and tutorials, is freely available athttps://plasmogenepi.github.io/PMO_Docs/. Key software components and datasets have archived releases with DOIs to ensure permanence, detailed in the Supplementary Text 1-5.

*Contact:*

## Introduction

Targeted sequencing is a powerful technique to directly address questions that do not require entire genomes (Bewicke-Copley *et al*. 2019). Targeted sequencing involves amplifying specific genomic regions, commonly using primer pairs, as in amplicon sequencing. The high sensitivity and specificity of this approach allow it to generate useful data even from samples containing low concentrations of the target organism, such as pathogens within their hosts, and have led to its adoption across a diverse range of applications that support disease response. Examples include human monkeypox virus outbreaks, pathogen identification and antimicrobial resistance marker surveillance, and wastewater surveillance of COVID-19 (Karthikeyan *et al*. 2022, Chen *et al*. 2023, Hernández-Neuta *et al*. 2023). Targeted sequencing can be rapidly deployed and adapted quickly as a response to emerging threats, including in low-resource settings where sequencing infrastructure may be limited (Ghansah *et al*. 2019). With data generation now widespread and continuing to increase, the lack of systematic and efficient analytical frameworks for these data has become a major factor limiting their impact.

A key advantage of targeted sequencing is the ability to directly obtain phased data from individual or paired-end reads within genomic areas of interest. This allows unambiguous linking of adjacent polymorphisms from nucleic acid fragments originating from the same input molecule, yielding microhaplotypes (Baetscher *et al*. 2018). Analysis of full microhaplotypes instead of independent SNPs are widely used in forensics and population studies as diverse multiallelic loci provide high resolution for evaluating genetic diversity and relatedness (Pakstis *et al*. 2012, Oldoni, Kidd, and Podini 2019, Hopken *et al*. 2023, Kwon and Shin 2024, Petrou *et al*. 2025, Thompson *et al*. 2025). It is important to note that individual variants such as SNPs and small indels can be derived from microhaplotypes, but microhaplotypes cannot be unambiguously inferred from individual variants, as phasing information is lost.

Microhaplotypes are becoming a widely employed tool for pathogen research and surveillance, particularly in organisms with complex infections (Tessema *et al*. 2022, Dash and Mallick 2024). A relevant example is *Plasmodium*, a genus of single-celled eukaryotic parasites, a subset of which cause human malaria. *Plasmodium*, which is haploid while within human hosts, infections are frequently polyclonal (i.e., multiple genetically distinct organisms are contained within a single sample), further elevating the value of retaining phasing information for multi-SNP microhaplotypes. Considerable community effort has been directed toward deriving analysis methods that take advantage of *Plasmodium* microhaplotype data; however, the development of reusable workflows to address key use cases requires the establishment of a standardised data storage format for microhaplotypes and associated metadata (Gerlovina *et al*. 2022, Murphy and Greenhouse 2024, Kleinecke *et al*. 2025, Ruybal-Pesántez *et al*. 2025). Challenges surrounding the lack of a standardised approach to representing these data not only extend to other eukaryotic pathogens, such as *Schistosoma, Leishmania*, and *Trypanosoma cruzi*, but also bacterial pathogens (Hay *et al*. 2017, Williamson *et al*. 2024, Furstenau *et al*. 2025). Establishing a robust data standard would therefore greatly facilitate consistent and interoperable analyses across diverse biological systems where microhaplotype data provide valuable insights.

Most established data formats were developed before use of microhaplotype data became common practice and were therefore not designed to handle these data efficiently. The variant call format (VCF) was designed primarily to convey individual variants within whole genome data and does not handle phased, multiallelic microhaplotypes well (Danecek *et al*. 2011). While imposing microhaplotype data into a VCF is possible, the resultant file would either include decomposed microhaplotypes spanning multiple records or include long, mixed length multi-allelic entries, where each possible haplotype is listed as an alternative allele. The first option would result in ambiguous representation or complete loss of phasing information for polyclonal samples. The second would result in incompatibility with downstream analysis methods built around this format, such as VCFtools, as it breaks the core assumption that each record is a single-site variant. The Biological Observation Matrix (BIOM) format and ESS-DIVE reporting formats can efficiently represent phased sequences from a single locus (McDonald *et al*. 2012, Velliquette *et al*. 2021). However, they were not designed to represent multi-allelic phased variation for multiple loci and lack the capacity to efficiently store these data.

The absence of an established standard for microhaplotype data has led to different groups creating bespoke, disparate solutions for data storage. This fragmentation hampers data sharing within laboratories, among collaborators, and across the wider community, leaving many datasets underutilized and requiring substantial manipulation for integration. Seemingly simple tasks like identifying metadata or defining a targeted sequencing panel are inconsistent, leading to errors and hindering comparison between datasets. Moreover, the lack of defined minimum information that should be shared to allow for analysis results in incomplete and fragmented datasets. Collectively, these issues have hindered data reuse, coordinated development of robust downstream analysis tools, and establishment of centralized repositories, leading to parallel efforts spent creating redundant software based on discordant formats.

To address the need for an established data standard for targeted sequencing data and associated metadata, the *Plasmodium* Genomic Epidemiology (PlasmoGenEpi) network has regularly convened and met with the wider community over the last four years to define a structured ontology for these data. The Portable Microhaplotype Object (PMO) outlined here integrates metadata and genomic data, allowing for efficient storage. The accompanying set of tools for file creation, manipulation, and sharing was designed to lower the barrier of entry, accelerating adoption and empowering the development of interoperable and reusable downstream analysis methods.

## Methods

### User-centered design

We sought to develop data standards and associated convenience utilities that would maximize accessibility and add value for a variety of potential users. Therefore, for the first two years of development, regular meetings were held with a focus group consisting of labs representing six universities (UCSF, UNC, Brown, UMass, John Hopkins, and Harvard). These groups collectively represent extensive experience working with diverse targeted sequencing panels and bioinformatic pipelines and have previously collaborated on data harmonization efforts, including a comparative MIP–Paragon analyses (Katairo *et al*. 2025a). This experience helped inform and motivate the development of the PMO framework. Through individual and group discussions, we identified consistent challenges in organizing amplicon sequencing data and metadata, leading to partial solutions ranging from inflexible, linear pipelines to manual storage of data in scattered spreadsheets. We identified a convergence point for data harmonization at the stage following basic bioinformatic allele calling, when samples have a set of assigned microhaplotype sequences and corresponding read counts (Figure 1). At this stage, data are the most comparable between labs (even when generated using different laboratory and bioinformatic methods), full microhaplotype sequence information is retained, and data storage requirements are lower than for raw sequence data. Thus, harmonizing data at this step for storage and data sharing offers the greatest flexibility for downstream analysis, providing disaggregated data to answer questions of interest.

**Figure 1.**
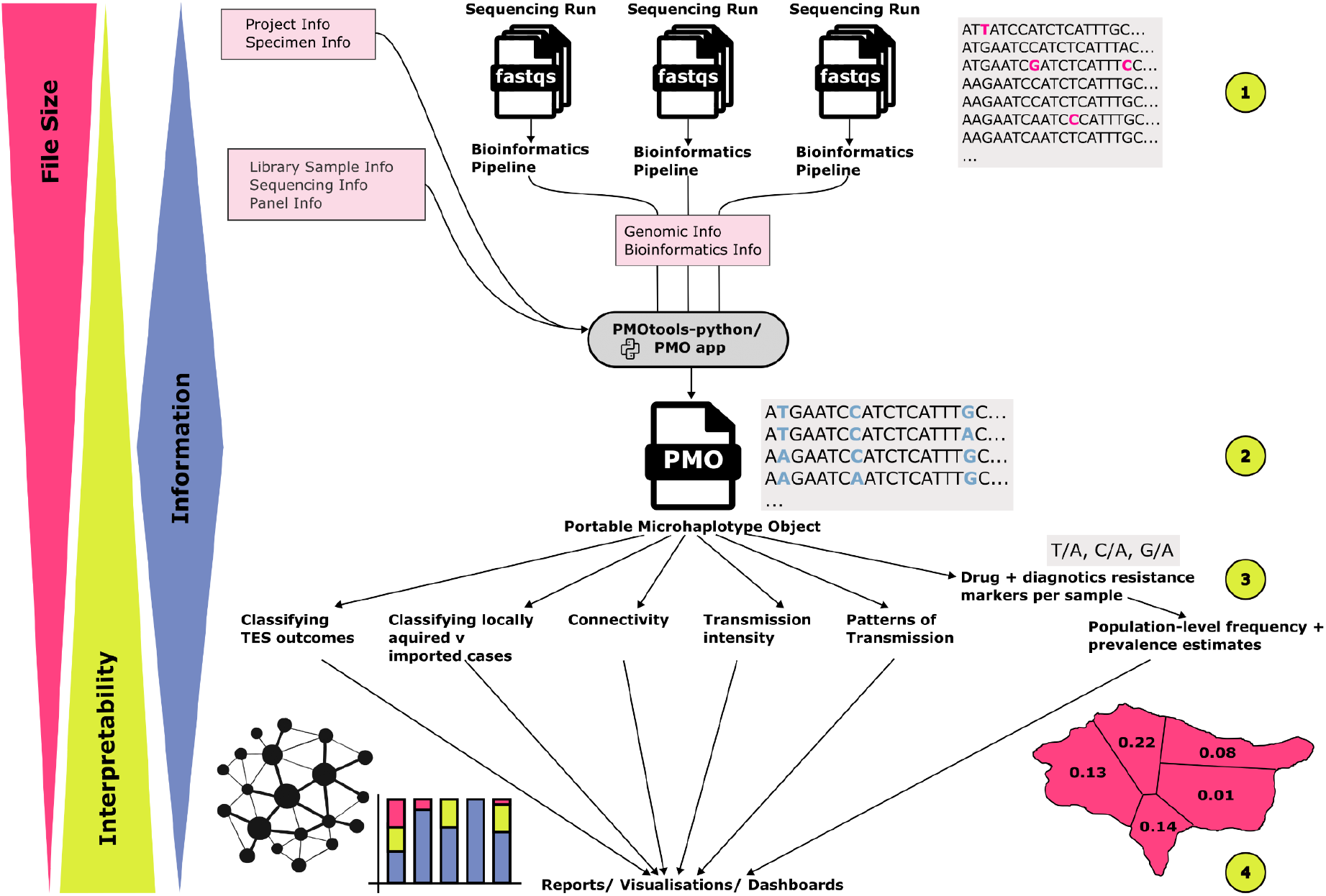
PMO as a convergence point (green circle 2) within the broader data ecosystem. This schematic outlines the flow of data in a typical workflow involving amplicon sequencing data. Green circles represent common stages for data sharing. Pink boxes indicate points at which information necessary for a PMO becomes available. (1) Raw sequencing data are generated, possibly from multiple sequencing runs at different points in time. FASTQ files for each sample represent a raw form of the data, with large files that are difficult to interpret without knowledge of the specific data-generating process or an appropriate allele-calling pipeline. At this stage, data are mostly shared with bioinformaticians and data repositories. Bioinformatics pipelines often require data from different sequencing runs to be processed separately to isolate any batch effects. After alleles are called, it is common to merge microhaplotype data from different runs. Harmonizing sources of data into a PMO file at this point allows an ideal convergence point for downstream analyses within the group, with collaborators, or with the broader community depending on the extent of data sharing. Simplified data for specific analyses, such as SNPs generated from microhaplotypes per sample or aggregated metrics such as allele frequency, can be easily derived from PMO. However, sharing data at this stage limits the scope of analyses that can be subsequently performed. The analysis examples presented are specific to *Plasmodium* molecular data use cases; however, the underlying principle of empowering downstream analysis applies broadly. (4) Interpreted results are shared e.g., through reports, manuscripts, and dashboards including maps, plots, and summary statistics. It is useful for information at this stage to include interpretation and simple representation. Though beyond the scope of this manuscript, establishing standards for downstream steps such as (3) and (4) may allow for integration of data and harmonization of analysis at additional stages of the workflow.

Linking metadata to sequencing data was also identified as a consistent challenge for data generators and users. These data were often stored separately in inconsistent formats and curated by different people, creating unnecessarily time-consuming and error-prone data merging steps. We therefore proposed an adaptable structure for combining data into one harmonized object at the stage after basic bioinformatics processing, empowering downstream analysis. A modular design was the most appealing to users, allowing flexibility to curate separate data components on an ongoing basis if necessary. For example, sample metadata could be recorded prior to sequencing then genomic data added as they are generated. This design allows for data to be continuously integrated into data storage systems and downstream analysis pipelines as they become available, allowing for ease of automation and improving reproducibility.

Once a core set of fields and concepts had been established, we engaged the broader community through small, informal discussion sessions at international meetings, including American Society of Tropical Medicine and Hygiene (ASTMH 2024) Annual Meeting, Genomic Epidemiology of Malaria (GEM 2024) Conference, and a meeting on malaria molecular surveillance standards hosted in Kampala, Uganda, in June 2025 by PHA4GE and the Africa Pathogen Genomics Initiative (Africa PGI) of the Africa Centres for Disease Control and Prevention (Africa CDC). Feedback from these sessions helped refine the ontology and ensure its relevance across a range of use cases and geographic settings. At these meetings, we addressed questions as they arose and encouraged participants to provide additional feedback through an open online comment form. Collectively, these engagement activities included approximately 50 participants from more than 30 institutions, including representatives from nine low- and middle-income countries (LMICs). Feedback collected through these channels was incorporated into successive revisions of the ontology. We subsequently presented the product at the ASTMH 2025 Annual Meeting as part of a scientific symposium and the PHA4GE Conference & IPSN Global Partners Forum 2025, providing a further opportunity to engage with the broader community and refine the framework prior to publication.

### PMO Ontology and File Format

The PMO version 1.1.0 stores genomic data and associated metadata in a single JavaScript Object Notation (JSON) formatted file (Supplementary Text 1, with detailed explanation of fields found at https://plasmogenepi.github.io/PMO_Docs/format/FormatOverview.html Supplementary Text 2). JSON allows for a relational structure and compact storage, is widely supported by computing languages, and is human readable. Having the ability to store all necessary data in a single file mitigates the frequent issue of genomic data becoming orphaned from metadata, allowing for clear versioning, easy data curation, and streamlined automation of data sharing including submission to repositories, downstream analysis, and visualisation. We used LinkML, the Linked Data Modeling Language, to define, document, and validate the relational structure of the data (linkml/linkml: Linked Open Data Modeling Language, n.d.). LinkML is increasingly adopted in the biological community to define data standards that are compliant with the FAIR principles of findability, accessibility, interoperability, and reusability (Wilkinson *et al*. 2016). Schema definition, as defined by LinkML, can be found here: https://plasmogenepi.github.io/portable-microhaplotype-object/. The LinkML-defined schema specifies the data structure of PMO, with various validation files (including JSONschema, SQLschema, JSON-LD, and OWL), and we have chosen the JSON file format as the default representation of PMO.

Fields are defined as either required or optional. Required fields define the minimum set of information needed for the data to be useful for downstream analysis. Optional fields include a larger set of elements that are used in a context-dependent manner, *i*.*e*., they may be useful or required for some analyses and may be included at the user’s discretion (*e*.*g*., granular details on location, method of sampling, or parasite density). Including a menu of commonly used optional fields allows for consistency in representation and may facilitate data organization for users while allowing flexibility in the extent of data shared. For example, a user may include details on the location of sampling for internal use (*e*.*g*., village) while opting to share less precise location data with collaborators or on public repositories. Due to the flexible nature of the JSON format, additional fields can also be incorporated by any user, however we recommend this be limited to fields that will not be widely reused by the community. Newly identified fields of broader utility should instead be defined and integrated into future versions of PMO.

To reduce redundancy and improve flexibility, the data are separated into modular tables that can easily be split downstream and integrated with other relevant ontologies. Figure 2A presents a schematic of the top-level tables, with their descriptions and representative examples provided in Table 1. Tables are unambiguously linked with internal indices. In the JSON file, the metadata tables are represented by a list of named entries for easy lookup per entry, while the microhaplotype data is encoded per sample per target with lookup values in the representative microhaplotype table. These sections can easily be exported into tabular format by pmotools-python. Where possible, fields were drawn from existing ontologies (e.g. the Minimum Information about any (x) Sequence (MIxS) standard from the Genomics Standards Consortium and SRA standards), with all required fields and over 60% of total fields derived from established standards (Yilmaz *et al*. 2011). Mapping of the source of fields can be found within the documentation (Supplementary Text 2, https://plasmogenepi.github.io/PMO_Docs/format/DevelopmentOfFormat.html).

**Table 1.** Details of top-level tables within the PMO schema. For each table, a short description and example fields are included. Optional fields/ tables are highlighted in grey. *The tables of library_sample_info and specimen_info only require identifiers but no additional metadata is required. A detailed description of all fields can be found within Supplementary Texts 1 (live athttps://plasmogenepi.github.io/portable-microhaplotype-object/) and 2 (live at https://plasmogenepi.github.io/PMO_Docs/format/FormatOverviewAdvanced.html).

| PMO table name | Description | Example fields |
| --- | --- | --- |
| **Required** |  |  |
| *pmo_header* | PMO-related information for this file. | *pmo_version*, *creation_date* |
| *representative_microhaplotypes* | Information on all of the microhaplotypes found within the population/ samples included in the PMO. | *seq*, *pseudocigar* |
| *detected_microhaplotypes* | For each *library_sample*, the microhaplotypes that were detected in that sample. | *mhap_id*, *reads* |
| *target_info* | Details of individual targets that make up panel(s) within PMO. | *Forward_primer_seq*, *reverse_primer_seq*, *insert_location*, *markers_of_interest* |
| panel_info | The collection of targets, optionally grouped by reaction, that were amplified and sequenced. | panel_name, target_id |
| **Optional** |  |  |
| targeted_genomes | Details on the genomes to which targets refer. | genome_version, taxon_id, url |
| specimen_info* | Details on the individual specimen collected. | specimen_name, specimen_taxon_id, collection_date, collection_country, host_age, host_sex, specimen_collect_device |
| library_sample_info* | Details of the specific amplification and sequencing of an individual specimen. | library_sample_name, library_prep_plate_info, run_accession |
| project_info | Information about the project(s) stored in this PMO. | project_name, project_description, project_contributors |
| sequencing_info | Details of the assay for the batch of samples sequenced together. | seq_platform, seq_instrument_model, seq_date |
| bioinformatics_methods_info | Details of the bioinformatics pipeline/methods used to generate the microhaplotypes. | program_version, program, program_description |
| bioinformatics_run_info | Runtime information for the batches processed by a bioinformatics method. | bioinformatics_run_name, run_date |
| read_counts_by_stage | The read counts per sample at different stages within the bioinformatics method/ pipeline, such as after demultiplexing or after denoising. | stage, read_count |

**Figure 2.**
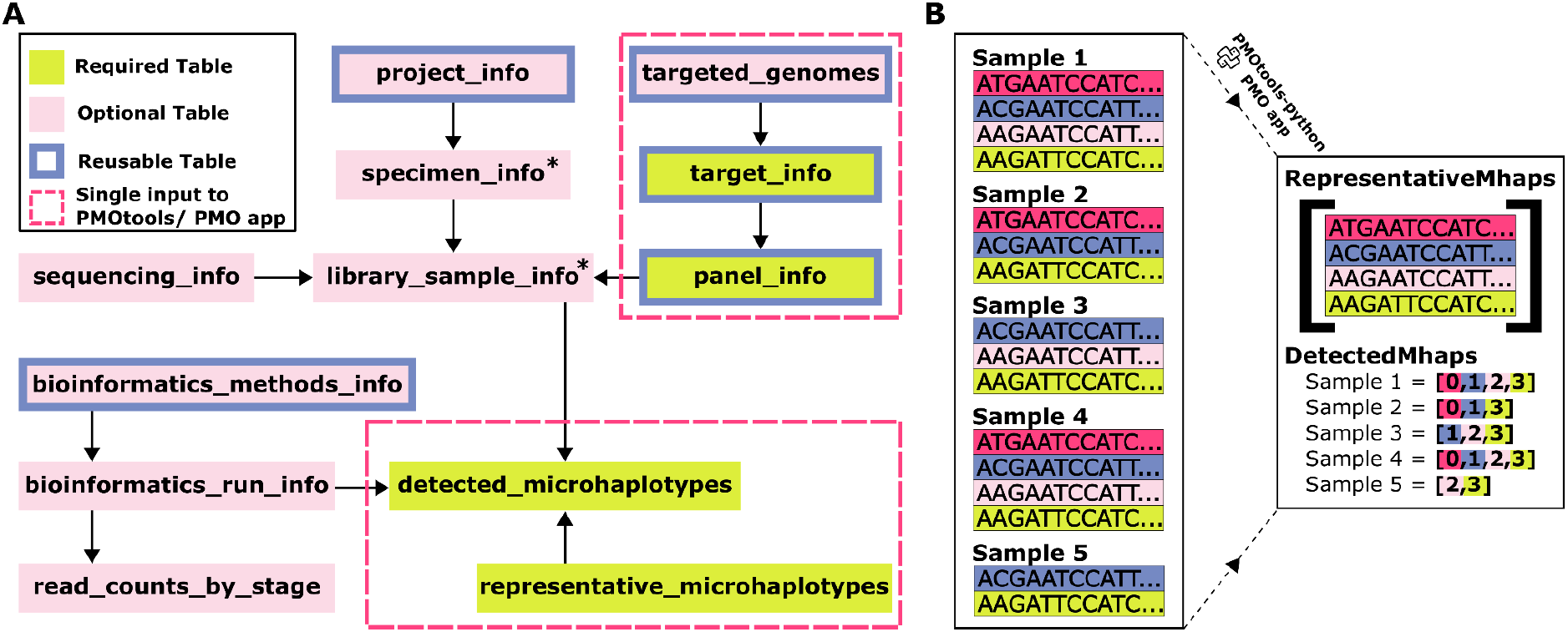
(A) Schematic of the top-level tables within PMO. Green and light pink boxes highlight required and optional tables respectively. Boxes with a blue border indicate tables that will commonly be reused across datasets. Pink dashed boxes highlight tables that are input collectively into pmotools-python or the PMO app via a single input table. *Tables specimen_info and library_sample_info require only an identifier, with additional information about the samples being optional. These identifiers can optionally be autogenerated through pmotools/pmo-app. (B) Illustration comparing current approaches to microhaplotype storage with storage within PMO. Current storage solutions often rely on long-form microhaplotype storage, with repeated listing of full nucleotide sequences, as shown on the left of panel B. In contrast, PMO replaces this with two efficiently linked tables, eliminating redundancy, as shown on the right of panel B.

A significant advance over available data formats provided by PMO is efficient, lossless storage of microhaplotype data, using two linked tables (Figure 2B). The full nucleotide sequence of each microhaplotype allele in the given dataset is represented only once in the file, in the *representative_microhaplotypes* table. Extra information can optionally be stored with these unique microhaplotype sequences including an abbreviated pseudocigar representation (a succinct representation of a microhaplotype sequence utilized by microhaplotype calling pipelines (LaVerriere *et al*. 2022)) and additional annotations. The microhaplotype alleles detected within each sample for each target are indicated using the index from the *representative_microhaplotypes* table, limiting repetition and allowing for efficient lookup.

Details for targeted sequencing panels are stored utilising a similar structure. Panels are often designed iteratively and contain re-used targets, or users may choose to sequence or analyze a subset of a full panel. To allow for flexibility in panel composition without needing to store details for each target multiple times, individual targets are defined within the *target_info* table, which includes primer sequences. Optionally, the intended genomic location of the forward primer, reverse primer, and insert can be defined using the genomic location fields. Panels are defined in the *panel_info* table by listing the indices of the targets that make up the panel. This structure allows for multiple related panels to be defined efficiently.

### pmotools-python

pmotools-python version 1.1.0 is an open-source Python package designed for converting data into the PMO format, performing basic data manipulation, and exporting information into other common formats (https://github.com/PlasmoGenEpi/pmotools-python, Supplementary Text 3). The package is pip-installable via PyPi and includes Python functions and a command-line interface that can be easily integrated into bioinformatics pipelines and analysis workflows. Two core inputs are required to build a PMO using pmotools-python: microhaplotype and panel information. Additional metadata describing the project, specimens, experiments, sequencing, and bioinformatics methods may also be included when available. For users unfamiliar with coding and Python, the PMO app (described below) offers a more accessible alternative for generating a PMO.

Many of these inputs can be created once and reused across groups, projects, and PMO files, since panels, bioinformatic pipelines, reference genomes, and sequencing methods are often replicated. In some cases, pmotools-python generates more than one of the tables within PMO from a single input. For example, a single dataframe containing genomic information can be processed using pmotools-python to generate the *representative_microhaplotypes* and *detected_microhaplotypes* tables, as depicted in Figure 2B. The functions include default column names for required fields that correspond to the PMO ontology. However, these column names can be set to align with users’ existing naming conventions. This empowers the automated conversion of data from established pipelines into PMO without the need to rename columns that users may be familiar with.

Functionality is also included to validate PMO compliance with the schema, perform basic data checks (*e*.*g*., no inappropriate duplication or missing metadata), subset PMO data (*e*.*g*., by date range or location), export to common outputs like allele tables, export all metadata to delimited plain text files or one Excel document, and provide basic summary statistics.

### PMO App

An open-source app with a graphical user interface was developed to provide a simple means for accurately creating a PMO without requiring coding or manual, error-prone data reshaping such as cutting and pasting (https://pmotools.app/, https://github.com/PlasmoGenEpi/pmotools-app, Supplementary Text 4). The app runs entirely on the user’s local machine using a standard web browser, requiring no additional software installation or coding knowledge. Users import tabular files by browsing or dragging and dropping them into the interface. The app then performs smart field matching to streamline data harmonization, matching disparate field names used across institutions into standardized required and optional fields within the PMO ontology. The user can quickly review assignments and, if needed, dynamically reassign them through drop down menus. Users may also add additional custom fields, if desired. Importantly, the app also allows for common sections to be saved. For example, information for specific genotyping panels and bioinformatic methods can be created once and easily reused for future data sets.

We also provide a multi-sheet Excel template that includes required and optional fields, along with descriptions of each PMO field. Although the tools are designed to accommodate diverse input formats, the template can help users get started with PMO creation. The template is available through https://pmotools.app and https://plasmogenepi.github.io/PMO_Docs. Full details on the format and the accompanying software can be found at https://plasmogenepi.github.io/PMO_Docs, which is continuously updated.

### Development and Maintenance of PMO

To ensure the continued development and maintenance of the format and its associated software, an active group of developers is working on the format and its implementation. Development is guided by regular discussions with a representative group of collaborating laboratories, as well as ongoing feedback from the broader user community. The development process is open and community-driven, and we welcome contributions from anyone interested in getting involved. Community members can participate by providing feedback, raising issues, suggesting enhancements, or submitting pull requests through the project GitHub repositories. Those interested in contributing are also encouraged to reach out directly to the development team to discuss ideas and opportunities for involvement.

## Results

We include 5 example datasets, demonstrating the versatility of PMO, including the ability to efficiently store data from large, multicenter projects, include data from different species, and easily organize different types of sequencing data for the same sample. See Supplementary Text 5 for more details, including the number of samples and targets per dataset.

The first dataset includes public genomic surveillance data of *Plasmodium falciparum* from 4 countries: Eswatini, Namibia, South Africa, and Zambia (Aranda-Díaz, Mwanza *et al*. 2025, Eloff *et al*. 2025, Nhlengethwa *et al*. 2025, Raman *et al*. 2025). These data were generated using the MAD^4^HatTeR drug-resistant targets (89 targets) amplicon sequencing panel applied to over 2000 specimens collected between 2022 and 2024 (Aranda-Díaz, Neubauer Vickers *et al*. 2025). Supplementary Text 5 includes a Jupyter Notebook outlining how the raw data were converted into the PMO format. A subset of specimens (522) were sequenced in replicate (Figure 3A). Sequencing information and microhaplotypes for all of these replicates are all stored in the PMO, efficiently linking to the unique specimen information. Simple metrics, such as the number of samples meeting specific quality control thresholds per country, can easily be extracted from PMO using pmotools-python (Figure 3B). More advanced analyses can also be performed efficiently; for example, drug resistance SNP frequencies can be assessed. Because metadata and genomic data are stored together, populations can be flexibly redefined e.g., by country or by province (Figure 3C,D).

**Figure 3.**
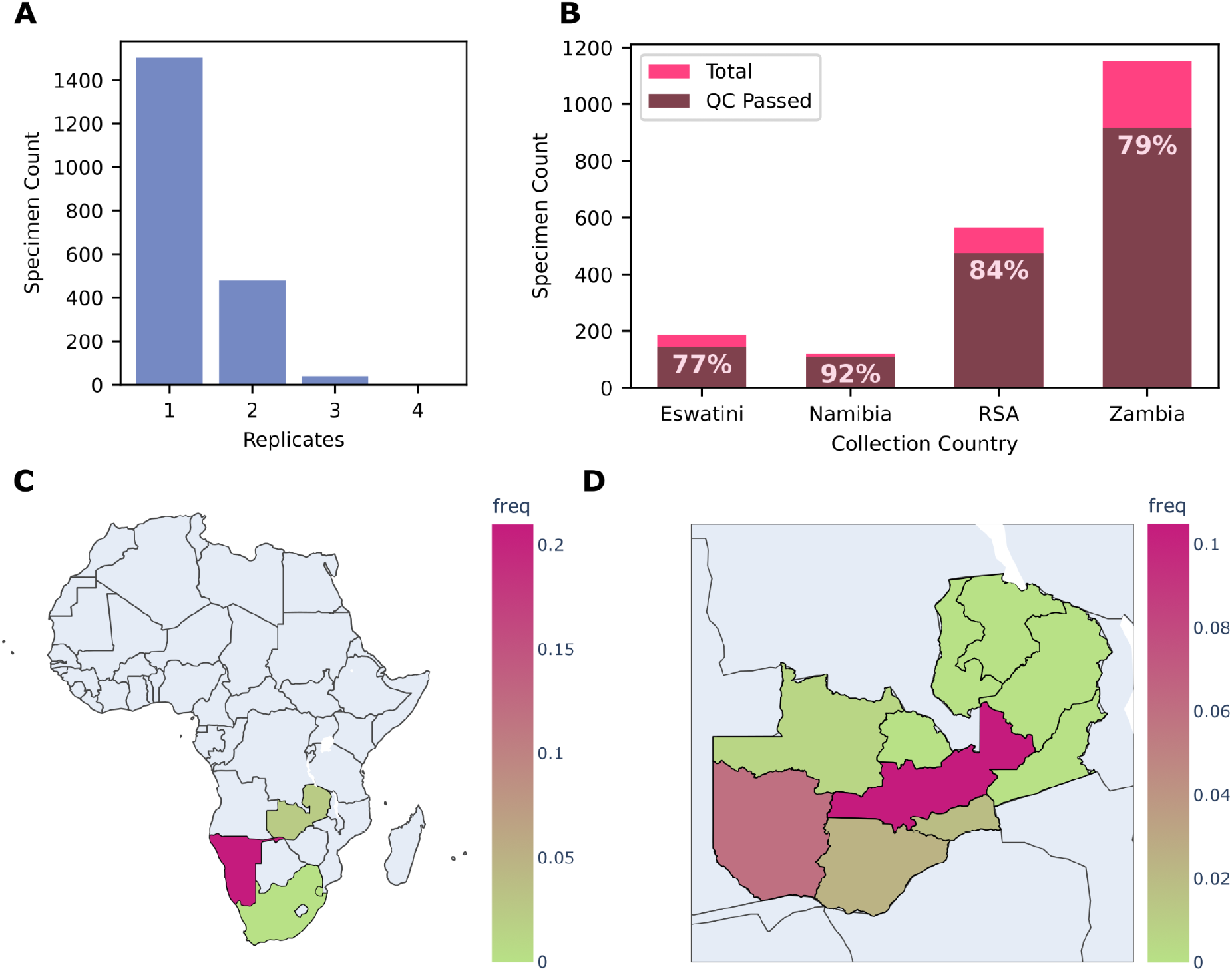
Reproduction of the analysis of genomic surveillance data of *Plasmodium falciparum* from 4 countries using PMO and pmotools-python. (A) Number of specimens sequenced in replicate. Duplicates for specimens can easily be dereplicated using PMOtools, in this case by retaining the *library_sample* with the highest total read count. (B) Number of specimens per country, with proportions flagged as passing QC filters (≥90% of targets with >50 reads). (C–D) Frequency of *k13* P441L across populations. By integrating metadata with genomic data, population definitions can be flexibly applied, for example by country (C) or by province in Zambia (D).

The second dataset includes *Anopheles* and *Plasmodium* data obtained from 4128 mosquito samples from Gabon that were sequenced as part of the ANOSPP (ANOpheles SPecies identification and Plasmodium detection) project (Makunin *et al*. 2022). The ANOSPP panel targets loci (64 targets) to determine mosquito species, population structure, and identify whether *Plasmodium* parasites are present in the mosquito. The unified storage of vector and parasite data in PMO format demonstrates the versatility of the modular, relational data structure, in this case including data from multiple organisms derived from the same sample. The notebook included shows how data from the project analysis pipeline were converted into PMO. We highlight the inclusion of additional fields, such as *sampling_location_size* and *plasmodium_detection_status* in the *specimen_info* table and *library_sample_info* table, respectively. More generally, fields specific to an assay or organism not included in the current ontology can easily be added using pmotools-python.

Next, we include a dataset of over 100 samples that were sequenced using two different *Plasmodium falciparum* assays - MAD^4^HatTeR amplicon sequencing (272 targets) and the DR23K molecular inversion probe (MIP) assay (118 targets) (Aranda-Díaz, Neubauer Vickers *et al*. 2025, Brhane *et al*. 2025, Katairo *et al*. 2025b). The modular structure of PMO avoids the replication of metadata for the specimens while retaining important assay-specific information such as the bioinformatic methods used. Importantly, results output from both assays can be directly compared. Notably, the MAD^4^HatTer panel and bioinformatic pipeline used here were identical to those employed for the first dataset described above, allowing reuse of the following PMO tables: *panel_info, targeted_genomes, target_info*, and *bioinformatics_method_info*.

Finally, we include two publicly available datasets from *Escherichia coli* (13 targets, 269 samples) and *Staphylococcus aureus* (27 targets, 164 samples) (Williamson *et al*. 2024, Furstenau *et al*. 2025). We downloaded metadata and genomic data directly from SRA. Several specimens had both targeted and whole-genome sequencing (WGS) data available. We generated microhaplotypes from the targeted data using SeekDeep and from the WGS data by first assembling with shovill via Bactopia and then extracting the corresponding targeted regions (Hathaway *et al*. 2018, Petit and Read 2020). We store all data in the same PMO, linking microhaplotypes from WGS and targeted sequencing to each specimen. This arrangement lets the user directly assess concordance between WGS and amplicon data while efficiently managing shared sample metadata.

### Data Size/ Scalability

A key advantage of PMO’s relational structure is efficient data storage, achieving ∼5-6-fold file size reduction relative to tabular representation of metadata and microhaplotypes, and up to ∼80-fold with additional compression using standard tools (e.g., gzip). A benefit of up to 16-fold can be achieved using PMO even if the tabular data are already compressed. File sizes scale with the number of samples, and even for the largest datasets described above, the final gzip compressed files remain under 3 MB, making them easily shareable via email. Supplemental Figure 1 compares file sizes of raw input data with both compressed and uncompressed PMO files for each dataset. This illustrates the substantial reduction in file size achieved when consolidating raw input data from diverse formats, including collections of Excel spreadsheets and CSV files, into PMO.

## Discussion

We developed PMO to address a critical gap in how microhaplotype data from targeted amplicon sequencing are stored and shared, with an initial implementation focused on *Plasmodium*, the pathogen responsible for malaria. Community input was prioritised throughout the 4-year development process, with regular working group meetings and broader engagement at conferences, ensuring diverse perspectives and expertise shaped PMO. By integrating metadata and genomic data efficiently in a single file, PMO enables consistent versioning as new data become available and protects against genomic data becoming orphaned from metadata. We developed two companion tools, pmotools-python and the PMO app, to improve accessibility and ensure seamless integration into automated pipelines. Given the modular structure of PMO, the framework should be generally applicable across the range of scientific endeavours being pursued with targeted amplicon sequencing.

PMO defines a minimum set of required fields and an extensive collection of optional ones, drawing from existing ontologies wherever possible. This ensures that data are interpretable by others when shared, enables downstream analysis, and secures the longevity of the data by eliminating reliance on users to reconstruct analysis steps or supply missing metadata. We leveraged JSON to allow for a relational structure of this complex data type, replacing memory intensive solutions with more efficient storage of data such as microhaplotypes. The LinkML framework generates various schema validators, including SQL validators, enabling the implementation and validation of the PMO schema in databases.

The full potential of this or any data standard in enabling data sharing and harmonization of analysis requires widespread adoption by those generating and consuming data. We have made initial efforts to facilitate broad uptake by providing extensive documentation, tutorials, training workshops, and easy-to-use tools. We designed pmotools-python and the PMO app to lower the barrier to implementation within current and future systems, and we encourage community participation in further expanding their functionality. These tools were designed to be approachable to both coders and users who would prefer an interactive interface. We are actively developing similar libraries in R and C++ to accommodate a wider range of users and plan to implement PMO interaction, e.g., dynamic data manipulation and visualization, into the app. Although pmotools-python includes functionality to export to several data formats, early adoption of PMO may require an initial investment to integrate with some existing downstream analysis tools. Centralized community resources can lower these barriers and are already being created. Efforts such as a website providing tutorials and tools landscaping (PGEforge), scripts that standardize inputs and outputs (PGEcore), benchmarking initiatives, and robust bioinformatics pipelines demonstrate how PMO is enabling coordinated development of such resources (Ruybal-Pesántez *et al*. 2025; *PlasmoGenEpi/PGEcore: Core Set of Scripts to Perform Common Downstream Analysis Functionalities for Plasmodium. Includes Wrappers of Tools and Bespoke Code to Perform Common Tasks*). Because these tools are being built around a common standard, they can be used and extended by anyone using PMO, making it easier for the community to build shared resources and amplifying the impact of new methods through broader uptake.

As this data format sits at a point of maximum information retention within the analysis workflow, we hope that the inheritance functionality of LinkML can be leveraged to reuse modules in future data standards that are established for outputs of downstream analyses. The ontology includes fields that we identified as relevant from existing ontologies. Our example datasets from bacteria, parasites, and vectors demonstrate that the format can already accommodate a wide range of use cases for storing and analyzing genomic data along with associated metadata. However, appropriate adaptation of domain-specific metadata fields will likely be required to maximize utility for organisms beyond *Plasmodium*. Fortunately, the flexible structure of PMO will easily accommodate this, and pmotools-python can still be used to create a PMO with any additional fields. In this case, we encourage the use of existing field names from ontologies such as the MIxS standard where possible, and users are invited to provide feedback of fields that could be valuable to include in future versions of PMO.

The PMO data format is explicitly designed for approaches that generate microhaplotypes spanning a consistent set of genomic positions, with all haplotypes sharing the same defined start and end coordinates (they cannot have “jagged” endings). As a result, the format is intended exclusively for this type of data and is not suitable for data that do not produce microhaplotype sequences as outputs. Consequently, PMO does not support data types such as microsatellite lengths, shotgun sequencing, or individual SNP genotypes. PMO is also only intended to store microhaplotype data and not the results of downstream analyses. For example, the PMO could be converted into SNP calls or used as input for population genomics (e.g., phylogenetic trees, admixture, within-/between-sample relatedness, allele frequencies, etc), but cannot and should not be used to store the results of those analyses. Finally, though there is no technical limit to the file size or amount of data stored in a PMO and we had no issue storing datasets with over 10,000 samples for 250 targets, the efficiency of the format will decrease at extreme sizes and care needs to be taken when exporting to formats such as Excel.

The work presented here offers an initial iteration of a solution to standardize storage of microhaplotype data generated via targeted sequencing, along with associated metadata. We intend this flexible, efficient means of data storage to facilitate organization of data for individual users, effective data sharing, and creation of appropriate repositories to maximize utility through reuse. By positioning PMO at the central point of the analysis workflow, where the maximum amount of information is retained, these standards also have the potential to catalyze nascent development of an interoperable ecosystem of easy-to-use analysis tools.

