## Supplementary Document for "The Portable Microhaplotype Object and Tools"

### Supplementary Materials

#### Supplementary Text 1 - PMO Schema

The LinkML-based schema is kept to date on GitHub. This repository has the yaml file that defines the schema and a live page that details the structure including all fields within PMO.

**GitHub (live version, continuously updated):**
https://github.com/PlasmoGenEpi/portable-microhaplotype-object
**Live Page:**
https://plasmogenepi.github.io/portable-microhaplotype-object/
**Zenodo Reference (version v1.1.0 from time of publication):**
https://doi.org/10.5281/zenodo.20550924

#### Supplementary Text 2 - PMO Documentation

The documentation for the PMO file format is kept up to date on its own GitHub page. This contains the details on the file format as well as tutorials and example files.

**GitHub (live version, continuously updated):**
https://github.com/PlasmoGenEpi/PMO_Docs
**Live Documentation Page:**
https://plasmogenepi.github.io/PMO_Docs/
**Zenodo Reference (version v1.1.0 from time of publication):**
https://doi.org/10.5281/zenodo.20551483

#### Supplementary Text 3 - PMO Python Implementation

**pmotools-python:** Toolkit for working with Portable Microhaplotype Objects (PMOs)
**GitHub (live version, continuously updated):**
https://github.com/PlasmoGenEpi/pmotools-python
Documentation describing installation, usage, and API examples is available and kept up to date at:
**Documentation (live, continuously updated):**
https://plasmogenepi.github.io/PMO_Docs/pmotools-python-usages/general_info.html
For reproducibility, version 1.1.0 of pmotools, used in this manuscript, has been archived at the time of publication:
**DOI (frozen at publication on Zenodo):** https://doi.org/10.5281/zenodo.20549057

#### Supplementary Text 4 - pmotools-app

**pmotools-app:** A simple web app built with Streamlit that helps users convert their data into the PMO format using the `pmotools` package.
**GitHub (live version, continuously updated):**
https://github.com/PlasmoGenEpi/pmotools-app
**Live app** is currently hosted here: https://pmotools.app/
**Documentation (live, continuously updated):**
https://plasmogenepi.github.io/PMO_Docs/pmotools-app-usage/pmotools-app.html
For reproducibility, version 1.1.0 of pmotools-app, used in this manuscript, has been archived at the time of publication:
**DOI (frozen at publication on Zenodo):** https://doi.org/10.5281/zenodo.20550287

#### Supplementary Text 5

All example datasets are available through the associated DOI (https://doi.org/10.5281/zenodo.20550920). The archive contains **five folders**, each corresponding to one dataset:

- **Dataset1:** Public genomic surveillance data of *Plasmodium falciparum* from four countries: Eswatini, Namibia, South Africa, and Zambia.
- **ANOSPP:** Combined *Anopheles* and *Plasmodium* data.
- **mips_v_mad4hatter:** Data from the MAD4HatTeR amplicon sequencing assay and the DR23K molecular inversion probe (MIP) assay comparison.
- **E_coli:** *Escherichia coli* datasets sourced from the Sequence Read Archive (SRA).
- **S_aureus:** *Staphylococcus aureus* datasets sourced from SRA.

For **Dataset1, S_aureus,** and **ANOSPP**, the archive includes all raw data files as well as Jupyter notebooks used to generate the PMO. Dataset1 additionally includes the notebook used to produce Figure 3. PMOs for **all five** datasets are included. The **S_aureus** dataset contains two PMOs: a PMO with the minimum amount required to build a PMO (microhaplotypes and primers) and a PMO that takes full advantage of all fields within the PMO schema.

Further details about the file structure and contents are provided in the accompanying README.md file with the archived data.

##### Supplementary Text 5 Table 1: Description of panels and number of samples in each of the datasets above.

| Dataset | Panel | Number of Targets | Total Unique Seqs | Average Length | Number of Library Samples |
|---|---|---|---|---|---|
| Dataset1 | MAD4HATTER-AB2 | 81 | 5967 | 191.8 | 2592 |
| ANOSPP | ANOSPP | 64 | 5648 | 204.5 | 4085 |
| mips_v_mad4hatter | MAD4HATTER | 272 | 2692 | 193.4 | 95 |
| mips_v_mad4hatter | MIPSDR23KE | 118 | 28139 | 191.5 | 196 |
| E_coi | coliseq | 13 | 271 | 148.7 | 269 |
| S_aureus | staph_aureus_Furstenau2025 | 27 | 143 | 128.7 | 164 |

#### Supplementary Text 6

**Glossary:**

**PMO** - Portable microhaplotype object

**VCF** - Variant Call Format - a text file format, with binary format options (BCF) for representing variant calls like insertions/deletions (INDELs) and single nucleotide polymorphisms (SNPs)

**BIOM** - Biological Observation Matrix - a JSON-based file format for representing observation-by-sample contingency tables with associated sample and observation metadata, commonly used in comparative omics and microbial ecology studies

**ESS-DIVE** - Environmental System Science Data Infrastructure for a Virtual Ecosystem - a U.S. DOE data archive for Earth and environmental science data, managing data, models, and software generated from research on terrestrial and subsurface environments

**LinkML** - Linked Data Modeling Language - an object-oriented data modeling framework that aims to simplify the production of FAIR, ontology-ready data, supporting schema specification and validation across formats like JSON-LD

**FAIR** - Findable, Accessible, Interoperable, and Reusable - a set of data guiding principles first proposed in 2016 to promote data management and stewardship practices that enhance data sharing and reuse, with an emphasis on machine-actionability

**SRA** - Sequence Read Archive - a bioinformatics database that provides a public repository for DNA sequencing data, especially short reads generated by high-throughput sequencing, run as a collaboration between NCBI, EBI, and DDBJ

**ENA** - European Nucleotide Archive - a repository providing free and unrestricted access to annotated DNA and RNA sequences, produced and maintained by the European Bioinformatics Institute and a member of the International Nucleotide Sequence Database Collaboration (INSDC)

### Supplementary Text 7

Table comparing features of each file format

| File Format | Handles phased data | Handles multiple targets | Stores Sample-level Metadata | Stores Sequencing/Bioinformatics Meta Data |
|---|---|---|---|---|
| PMO | Yes | Yes | Yes | Yes |
| VCF | No | No | No | No |
| Biome | Yes | Yes | Yes | No |
| ESS-Dive | Yes | No | Yes | Yes |

```json
{
  "pmo_header": {
    "pmo_version": "1.1.0",
    "creation_date": "2026-05-15",
    "generation_method": {
      "program_name": "pmotools-python",
      "program_version": "1.1.0"
    }
  },

  "library_sample_info": [
      {
      "library_sample_name": "SRR30825770",
      "specimen_id": 0,
      "panel_id": 0
    },
    ....
  ],
  "specimen_info": [
      {
      "specimen_name": "SRR30825770"
    },
    ....
  ],
  "panel_info": [
    {
      "panel_name": "staph_aureus_Furstenau2025",
      "reactions": [
        {
          "reaction_name": "full",
          "panel_targets": [
            0,
            1,
            .....
          ]
        }
      ]
    }
  ],
  "target_info":[
      {
      "target_name": "SA_131432",
      "forward_primer": {
        "seq": "GTCCAGGTAGCATGATT"
      },
      "reverse_primer": {
        "seq": "TGTCATACCAGTTAGGAATCACA"
      }
    },
    ....
  ],
  "representative_microhaplotypes":{
      "targets": [
      {
        "microhaplotypes": [
          {
            "seq": "CAATATAATAACCTAATAAAATGTTTAGGTCAACCTAAATTTATTTTAATTTTTTTAAAAGTAT
          },
        ],
        "target_id": 8
      },
      ....
      ]
  },
  "detected_microhaplotypes": [
      {
      "library_samples": [
        {
          "library_sample_id" : 1,
          "target_results": [
            {
              "mhaps_target_id": 0,
              "mhaps": [
                {
                  "mhap_id": 6,
                  "reads": 428
                }
              ]
            },
            ....
        ]
      }
      ....
    ]
    }
    ...
  ]
}
```

Supplementary Figure 1: A visual example of what the internal structure of the PMO file looks like. More details on these can be found at the PMO_Docs website at https://plasmogenepi.github.io/PMO_Docs/format/FormatOverviewAdvanced.html.

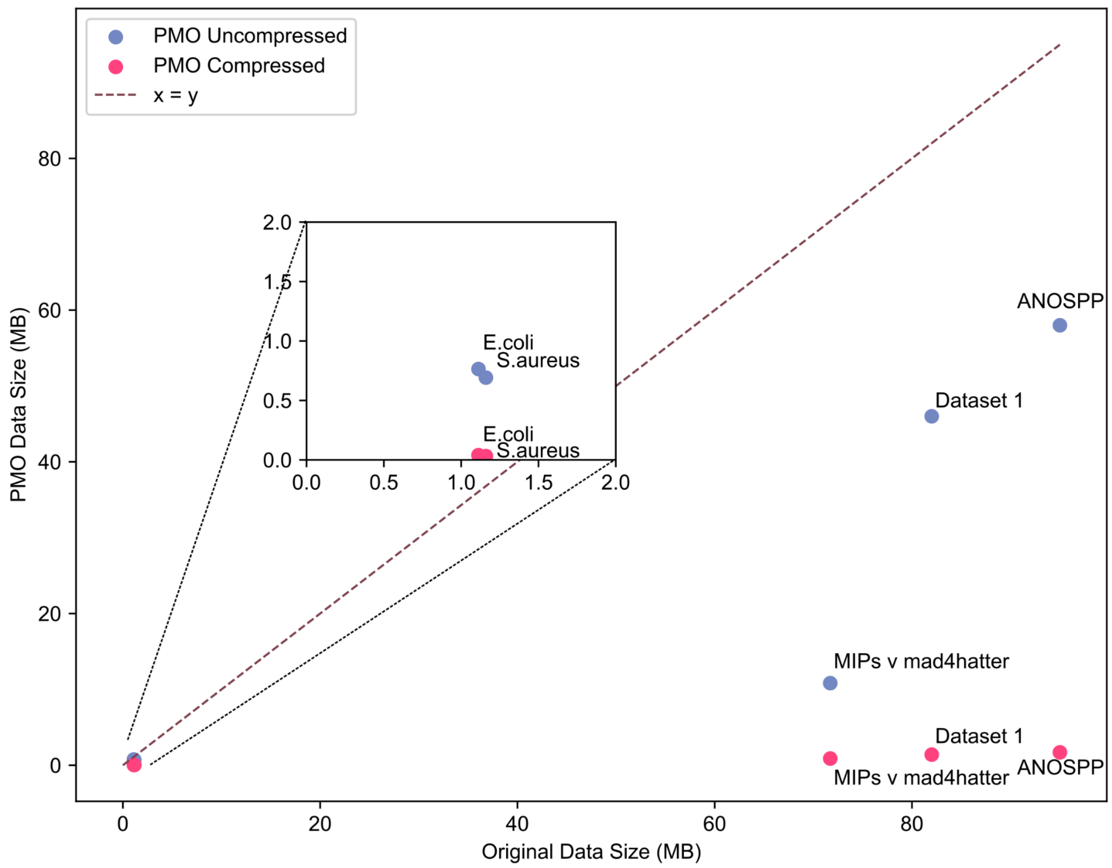


**Supplementary Figure 2. Comparison of original dataset size and size after conversion to PMO.** Scatter plot comparing the size of the original data (x-axis) and the corresponding PMO data (y-axis), for both uncompressed and compressed PMO outputs. All datasets show substantial reductions in size as a PMO. The inset axis provides a magnified view of the low-MB range (*S.aureus* and *E. coli*) to highlight the pronounced compression effects in small datasets. All values that were used to create the plot above can be found in the table below for comparison. All sizes are in megabytes. The sizes of the original data, the original data in standard gzip compression, the size of the PMO file, the size of the gzip PMO file, and the ratios of the PMO to the original data are shown.

| Dataset | Uncompressed Original Data Size (MB) | Compressed Original Data Size (MB) | PMO Data Size (MB) | Uncompressed PMO to Uncompressed Original Ratio | Compressed PMO Data Size (MB) | Compressed PMO to Compressed Original Ratio | Compressed PMO to Uncompressed Original Ratio |
|---|---|---|---|---|---|---|---|
| Dataset 1 | 69.234375 | 3.1484375 | 13.046875 | 5.306586826 | 1.09765625 | 2.868327402 | 63.0747331 |
| ANOSPP | 97.3046875 | 11.2109375 | 19.12890625 | 5.086787829 | 1.5234375 | 7.358974359 | 63.87179487 |
| MIPs v mad4hatter | 70.609375 | 14.11328125 | 10.5703125 | 6.679970436 | 0.86328125 | 16.34841629 | 81.7918552 |
| E.coli | 1.0859375 | 0.08203125 | 0.74609375 | 1.455497382 | 0.0390625 | 2.1 | 27.8 |
| S.aureus | 1.1328125 | 0.12890625 | 0.67578125 | 1.676300578 | 0.03125 | 4.125 | 36.25 |
